# The CALERIE™ Genomic Data Resource

**DOI:** 10.1101/2024.05.17.594714

**Authors:** CP Ryan, DL Corcoran, N Banskota, C Eckstein Indik, A Floratos, R Friedman, MS Kobor, VB Kraus, WE Kraus, JL MacIsaac, MC Orenduff, CF Pieper, JP White, L Ferrucci, S Horvath, KM Huffman, DW Belsky

## Abstract

Caloric restriction (CR) slows biological aging and prolongs healthy lifespan in model organisms. Findings from CALERIE-2™ – the first ever randomized, controlled trial of long-term CR in healthy, non-obese humans – broadly supports a similar pattern of effects in humans. To expand our understanding of the molecular pathways and biological processes underpinning CR effects in humans, we generated a series of genomic datasets from stored biospecimens collected from n=218 participants during the trial. These data constitute the first publicly-accessible genomic data resource for a randomized controlled trial of an intervention targeting the biology of aging. Datasets include whole-genome SNP genotypes, and three-timepoint-longitudinal DNA methylation, mRNA, and small RNA datasets generated from blood, skeletal muscle, and adipose tissue samples (total sample n=2327). The CALERIE Genomic Data Resource described in this article is available from the Aging Research Biobank. This multi-tissue, multi-omic, longitudinal data resource has great potential to advance translational geroscience.

## Introduction

Caloric restriction (CR) involves a reduction of caloric intake by 10-20% or more while maintaining adequate protein and micronutrient levels^1^. In laboratory experiments, CR slows biological aging and prolongs lifespan in multiple model organisms^2^. In humans, observational studies of individuals who self-impose CR^3,4^, short-term trials involving participants with obesity^5^, and involuntary CR among participants in the Biophere-II experiment^6^ broadly support some of the health and longevity promoting effects shown in model organisms^7^. The CALERIE-2™ (Comprehensive Assessment of Long-term Effects of Reducing Intake of Energy) study (NCT00427193 at clinicaltrials.gov), is the first-ever randomized controlled trial of CR in healthy, non-obese humans. CALERIE-2™ is a phase II, multicenter, randomized controlled trial (RCT) that tested the effects of a 24-month CR intervention in healthy non-obese adult men and women. Although the trial did not achieve durable modification of its primary endpoints, resting metabolic rate and core temperature^8^, analysis of participants’ clinical laboratory values broadly support the hypothesis that CR promotes healthy aging; participants randomized to the CR condition experienced broad improvements in their cardiometabolic health relative to those in the control “ad libitum” (AL) condition^9^ and exhibited a slowed pace of biological aging, as measured from algorithms based on clinical laboratory values^10,11^. These observations motivated efforts to elucidate underlying cellular-level mechanisms, including through projects funded by the National Institute on Aging at Columbia Univeristy, Duke University, and the University of California Los Angeles. The outcomes of these projects, including SNP genotype, DNA methylation, and messenger and small RNA sequencing data derived from blood, muscle, and adipose tissues collected from participants at pre-intervention baseline and at the 12- and 24-month follow-up assessments comprise the the CALERIE™ Genomic Data Resource.

### Description of the Trial

Extending CALERIE™ phase I^12^ in both scale and duration, CALERIE-2™ recruited a total of 220 subjects and assigned them in a 2:1 allocation to a CR treatment group or ad libitum (AL) control arm^13^. Subjects were randomly-assigned to CR or AL groups stratified on study site, sex, and body mass index. Participants in the CR group were assigned to a protocol designed to result in a 25% reduction in caloric intake relative to estimated energy requirements at enrollment. CR participants received an intensive behavioral intervention that included individual and group sessions, a meal provision phase, digital assistants to monitor caloric intake, and training in portion estimation and other nutrition and behavioral topics^14^. Adherence was assessed using measures of energy expenditure using the doubly-labelled water method as well as expected changes in body composition. The duration of the study for both CR and AL participants was 2-years.

Throughout the 2-year study duration, starting at baseline prior to randomization and recurring at months 1, 3, 6, 9, 12, 18, and 24, participants were evaluated for a range of pre-specified anthropometric, psychological, and physiological outcomes^8^. Blood samples were collected every six months. At baseline, 12-months, and 24-months, whole blood and samples were collected and banked. In addition, a subset of participants agreed to biopsies of adipose and muscle tissue at baseline, 12-months, and 24-months. From these samples SNP-based genotypes, DNA methylation, mRNA, and small RNAs were assayed. Here, we describe these datasets and provide an overview of this data resource (**Fig. 1**).

**Figure 1.**
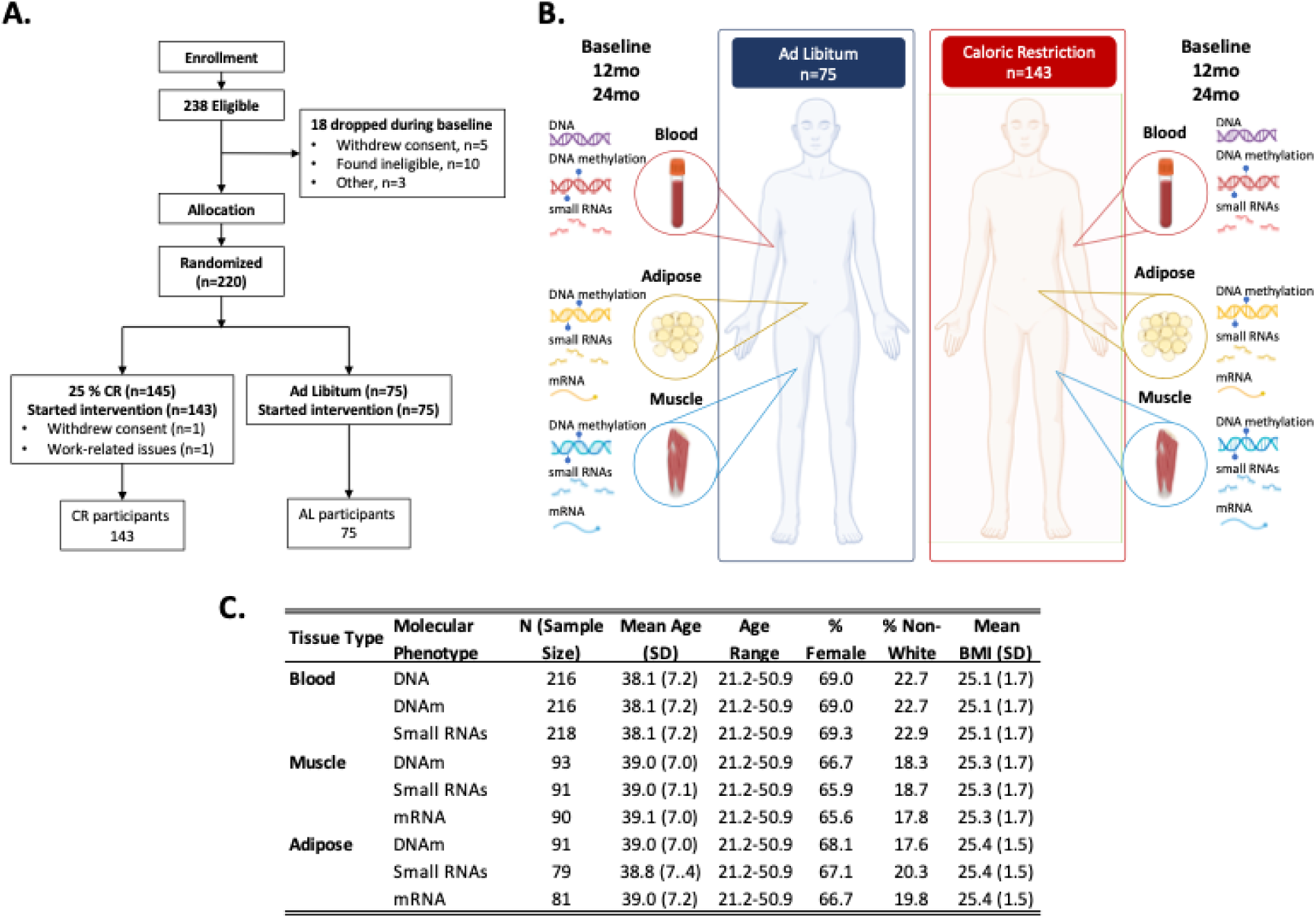
Study design, participant information, and overview of molecular datasets generated for the CALERIE™ caloric restriction trial. Panel A shows the Consort diagram showing enrollment, eligibility, and final sample sizes by treatment group. Panel B shows a schematic of molecular data types and tissue sources. Genetic material (DNA) was derived from blood samples for both ad libitum (AL) and caloric restriction (CR) groups at baseline. DNA methylation was derived from blood, muscle, and adipose samples for both AL and CR at baseline, 12-months, and 24-months. Small RNAs were derived from blood, muscle, and adipose samples for both AL and CR at baseline, 12-months, and 24-months. mRNA was derived from muscle and adipose samples for both AL and CR at baseline, 12-months, and 24-months. Panel C shows a table of demographic information for participants that contributed data to individual tissue level molecular datasets. Total sample size, mean and standard deviation in age, age range, percent female, percent who self-reported as other than white, and mean and standard deviation of BMI. Figure created using images from Biorender.com

More details about the CALERIE™ trial, including study protocols and on-going and published research, are available at https://calerie.duke.edu. Data can be accessed through the Aging Research Biobank (https://agingresearchbiobank.nia.nih.gov/studies/calerie/). Data use is restricted to non-commercial use in studies to determine factors that affect age-related conditions. Biospecimens are available, limited to research on effects that caloric restriction may have on aging and aging-related diseases.

## Results

### Overview

Of 238 eligible enrolled participants, n=220 were allocated to the trial and randomized to CR and AL treatment groups (**Fig. 1**). Detailed study procedures were published previously^8,12,15^ and are described here in brief. Two participants in the CR treatment group withdrew from the study, leaving n=143 participants in the CR group and n=75 in the AL group. In total, genomic data was produced for n=218 unique individuals. SNP data were available for 216 participants. DNAm and RNA sample coverage varied by treatment group, time-point, and tissue type (**Fig. 2 and Supplemental Figure S1**). A sample availability matrix for all samples by molecular data type, tissue, and timepoint is available in **Table S1**.

**Figure 2.**
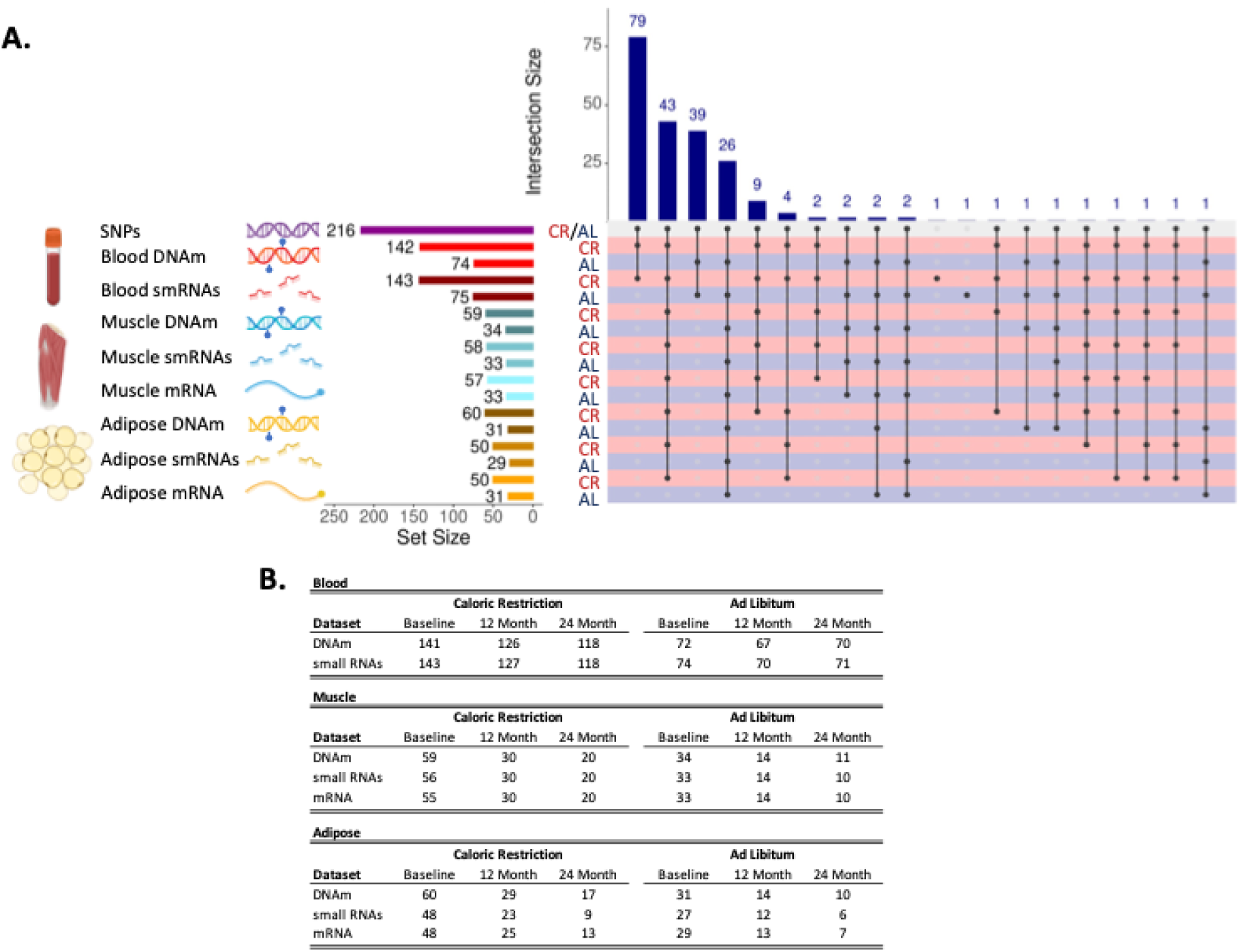
Panel A shows an upset plot showing overlap of available datasets across treatment group, tissues, molecular data type. The left-hand side shows tissue type (blood, muscle, or adipose), type of molecular data type (SNPs, DNAm, smRNAs, or mRNA), set size for each tissue and data type combination in number of unique individuals, and group (CR or AL). The bottom right-hand side shows points and connecting lines indicating overlapping intersections across tissues and data types color coded by treatment group (CR = red, AL = navy). The top right hand side shows a barchart indicating sample sizes for the overlapping intersections of tissue and datatypes. Each tissue and molecular data type combination is linked to a corresponding color scheme as follows: genomic variation (purple DNA), blood DNA methylation (red DNA with lollipops), blood small RNAs (smRNAs; dark red RNA fragments), muscle DNA methylation (blue DNA with lollipops), muscle smRNAs (blue RNA fragments), muscle mRNA (blue single RNA strand), adipose DNA methylation (yellow DNA with lollipops), adipose smRNAs (yellow RNA fragments), adipose mRNA (yellow single RNA strand). Panel B shows Individual sample numbers by molecular data type, treatment group (Caloric Restriction or Ad Libitum) and follow-up visit (Baseline, 12 Month, or 24 Month) across different tissue types. For some individuals, samples were available at follow-up but not baseline (or vice versa), so baseline numbers for a tissue and treatment group combination will not always match sample sizes in Figure 2 Panel A. Figure created using images from Biorender.com.

### Genotyping

Genotypic data is available for all but two participants who were randomized, provided consent, and completed the trial (n=216). **Figure 3 Panel A** illustrates genetic ancestry of CALERIE participants as reflected in the first two principal components of the genome-wide single-nucleotide polymorphism (SNP) data. **Figure 3 Panel B** illustrates the percent genetic variation explained by the top 10 genetic principal components.

**Figure 3.**
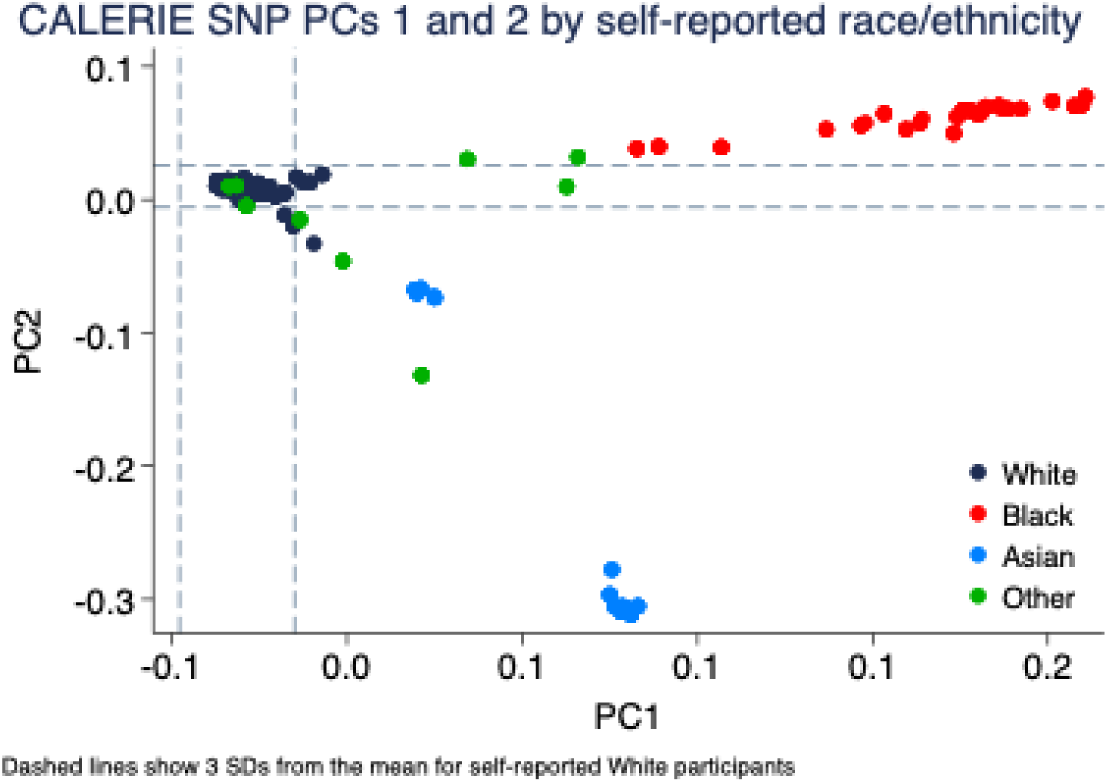
SNP-based principal components scores for CALERIE™ participants. The figure shows the top two SNP-based principal components (PCs) for CALERIE™ participants (n=217), colored by self-reported race/ethnicity. Dashed lines indicate three standard deviations from the mean for participants who self-reported race/ethnicity as White. The top-5 SNP-based principal-components explained 11.9% (PC1) 4.8% (PC2), 1.5% (PC3), 1.4% (PC4), and 1.3% (PC5) of the genetic variation among CALERIE participants.

### DNA methylation

DNA methylation (DNAm) was assayed from DNA extracted from whole blood, skeletal muscle, and adipose tissue using Illumina EPICv1 arrays (Fig. 1B). Assays of whole blood DNAm were completed by the Kobor Lab at the University of British Columbia as part of NIH grant R01AG061378. Assays of muscle and adipose DNAm were completed by the UCLA Neuroscience Genomics Core (UNGC) as part of NIH grant U01AG060908 using DNA extracted from banked tissue by the VB Kraus lab in the Duke Molecular Physiology Institute. Quality control and preprocessing for muscle and adipose tissue DNAm was carried out by the Geroscience Computational Core of the Robert N. Butler Columbia Aging Center.

Blood DNAm is available for at least one timepoint for n=216 participants (n=142 participants in the CR group and n=74 in the AL group)^16^. Sample sizes for blood DNAm for individual timepoints by treatment group are provided in **Fig. 2A**. Muscle DNAm is available at least one timepoint for n=93 (n=59 participants in the CR group and n=34 in the AL group). Sample sizes for muscle DNAm for individual timepoints by treatment group are provided in **Fig. 2A**. Adipose DNAm is available for at least one timepoint for n=91 participants (n=60 participants in the CR group and n=31 in the AL group). Sample sizes for adipose DNAm for individual timepoints by treatment group are provided in **Fig. 2A**. Muscle and adipose DNAm data have not been published previously.

To facilitate the rapid appraisal of the underlying data structure and missingness, we generated a series of summary datasets. For each tissue, we provide a set of tables listing the probes that passed quality control (**Supplemental Data Files 1-4, 9-12, 17-20**) and their beta-value distributions and missingness at each timepoint (**Supplemental Data Files 5-8, 13-16, 21-24**).

In addition to CpG methylation levels, we also computed a series of composite variables from the blood DNAm data. To help account for technical variability across samples, we provide the top principal components of EPIC-array control-probe beta values for blood, adipose, and muscle^17^. For contrast between tissues, we illustrate deconvoluted cell types (adipose, epithelial, fibroblast, and immune cells) for blood, muscle, and adipose using hierarchical in **Figure S2**.

We also include in this database the series of epigenetic clocks and estimated white blood cell proportions reported previously^18^ as well as a newly-derived set of estimated white blood cell proportions^19^. Intercorrelations of chronological age and five epigenetic clocks (PCHorvath, PCHannum, PCPhenoAge, PCGrimAge, and DunedinPACE) at the baseline assessment, prior to treatment, are provided in **Figure 4**. Intercorrelations of chronological age and ten epigenetic clocks (Horvatth, PCHorvath, Hannum, PCHannum, PhenoAge, PCPhenoAge, GrimAge, PCGrimAge, and DunedinPACE) at the baseline assessment, prior to treatment, are provided in **Figures S3 and S4**. As published elsewhere, the clocks show low to moderate intercorrelation with one another, reflecting the different methods of their construction^20,21^. Relative proportions of each of the 12 white blood cell types estimated at baseline measurement, prior to treatment, using previously described methods^19^ are graphed in **Figure 5**.

**Figure 4.**
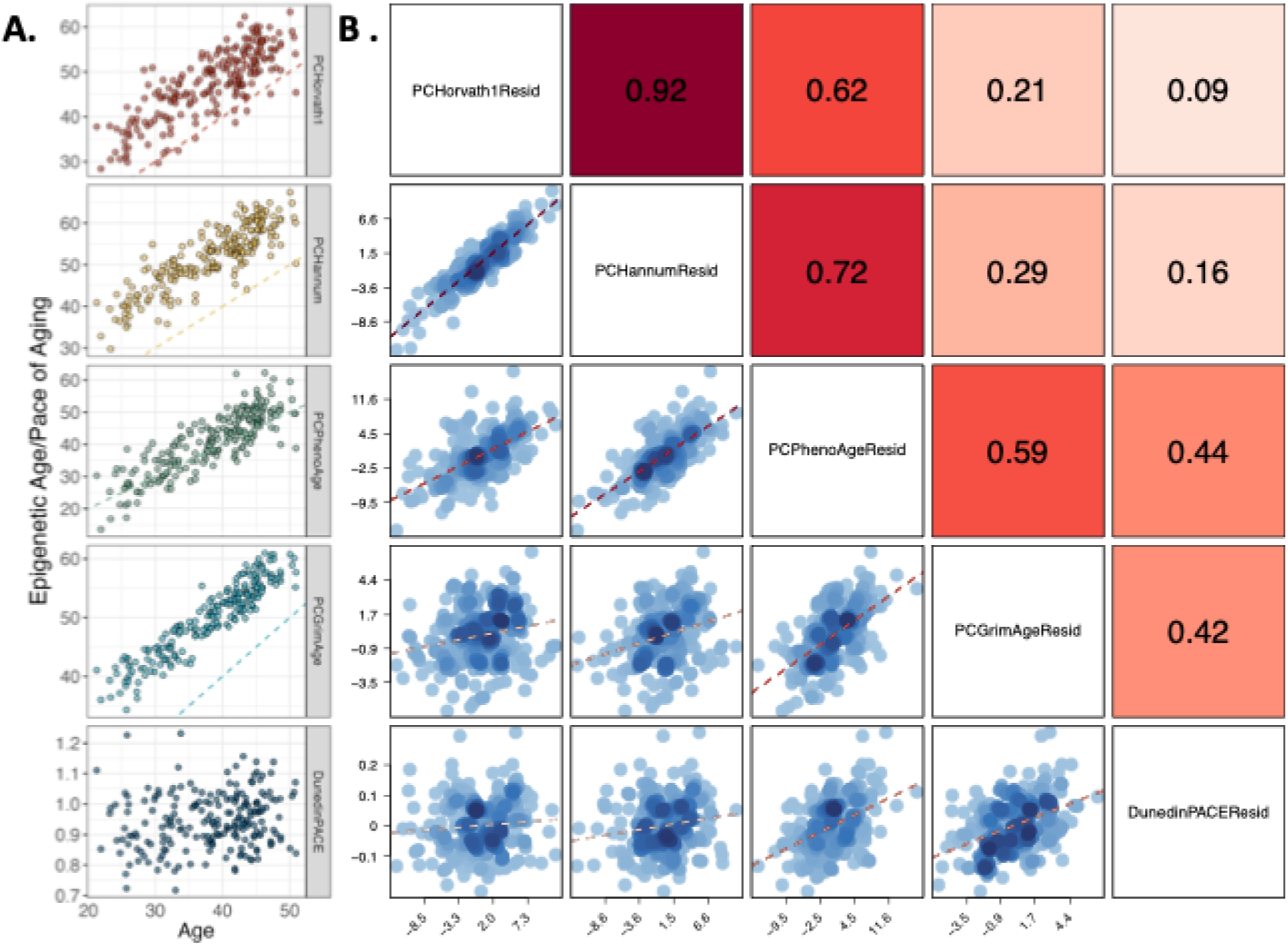
Associations of DNA methylation measures of aging with chronological age and age-residualized DNA methylation measures of aging with each other. Panel A shows DNA methylation measures of aging (Y-axis) against chronological age (X-axis) for n=212 men and women at pre-intervention baseline. The dashed colored line on each facet is the line of identity (intercept=0, slope=1), indicating where predicted epigenetic age or pace of aging would equal chronological age. Correlations with chronological age are as follows: PC Horvath Clock r=0.84, PC Hannum Clock r=0.88, PC PhenoAge Clock r=0.85, PC GrimAge Clock r=0.92, DunedinPACE r=0.15. Panel B shows correlations between age-residualized DNA methylation measures of aging for n=212 men and women at pre-intervention baseline with each other. The dashed red line on each facet is the fitted regression slopes. Pearson correlations between DNA methylation measures of aging are shown on the upper diagonal facets, with the shade of the facet indicating the strength of the correlation.

**Figure 5.**
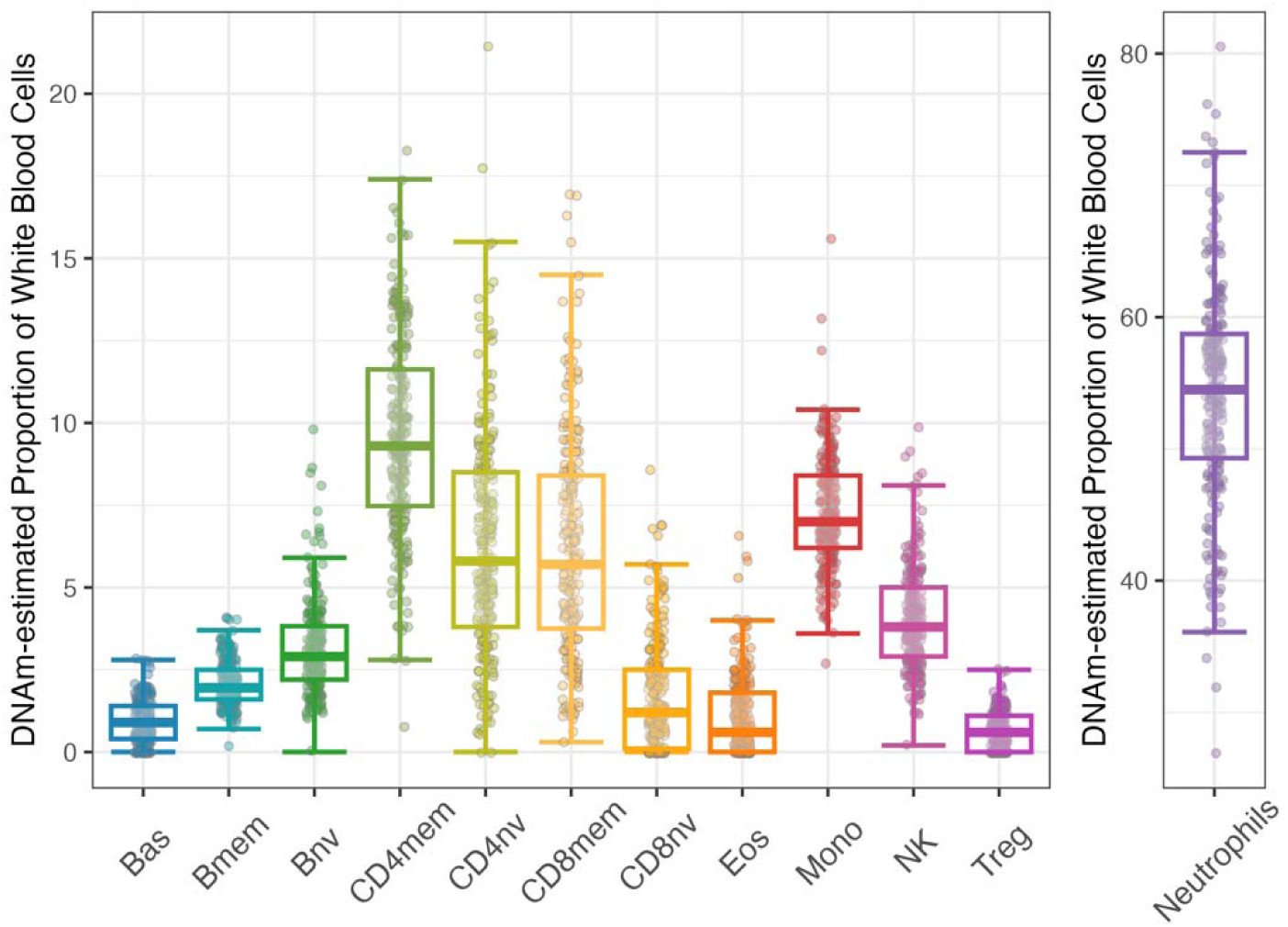
Relative proportions of 12 white blood cell types estimated for all participants at baseline. The figure shows the estimated relative proportions of 12 white blood cell types for all participants at baseline, prior to treatment (Bas=basophils, Bmem=memory B cells, Bnv=naïve B cells, CD4mem=memory CD4T cells, CD4nv=naïve CD4T cells, CD8mem=memory CD8T cells, CD8nv=naïve CD8T cells, Eos=Eosinophils, Mono=Monocytes, NK=Natural Killer cells, Treg=T regulatory cells, Neutrophils=Neutrophil cells). Individual points represent individual observations. Box and whisker plots showing median (horizontal line), 25^th^ quartile (bottom of the box), 75^th^ quartile (top of the box), and minimum and maximum values within 1.5 × IQR (interquartile range; top and bottom segments, respectively). Neutrophils are plotted on a separate scale.

### RNAseq

RNA sequencing was performed for three tissues: plasma, skeletal muscle, and adipose tissue (Fig. 1B). All RNA was extracted from plasma at the Duke Molecular Physiology Institute from stored plasma, skeletal muscle, and adipose-tissue biospecimens. Plasma and adipose RNA were sequenced at the Duke University Center for Genomic and Computational Biology. Muscle RNA was sequenced at the Center for Cancer Research Sequencing facility (National Cancer Institute). Messenger RNA (mRNA) and small RNA (smRNA) datasets were generated for skeletal muscle and adipose tissues. Only smRNAs were generated from plasma due to mRNA sample quality issues.

### Small RNAs

Plasma smRNAs were available for at least one timepoint for n=218 participants (n=143 in the CR group and n=75 in the AL group). Sample sizes for plasma smRNAs for individual timepoints by treatment group are provided in **Fig. 2A**. Muscle smRNAs are available for at least one timepoint for n=91 participants (n=58 in the CR group and n=33 participants in the AL group). Sample sizes for muscle smRNA for individual timepoints by treatment group are provided in **Fig. 2A**. Adipose smRNAs are available for at least one timepoint for n=79 participants (n=50 in the CR group and n=29 participants in the AL group). Sample sizes for adipose smRNAs for individual timepoints by treatment group are provided in **Fig. 2A**.

### Messenger RNA

Muscle mRNA is available for at least one timepoint for n=90 participants (n=57 in the CR group and n=33 in the AL group)^22^. Sample sizes for muscle mRNA for individual timepoints by treatment group are provided in **Fig. 2A**. Adipose mRNA is available at least one timepoint for n=81 participants (n=50 in the CR group and n=31 in the AL group). This dataset was generated from an independent and larger set of samples than what has been published previously^23^.

Sample sizes for this new adipose mRNA for individual timepoints by treatment group are provided in **Fig. 2A**. Adipose mRNA analysis has not been published previously. An overview of adipose mRNA response to CALERIE intervention in this dataset is reported in **Fig. 6**. Differential expression analysis identified 605 genes modified by CR at the 12-month follow-up (**Fig. 6A**; 309 upregulated and 296 downregulated) and 734 genes at the 24-month follow-up (**Fig. 6C**; 330 upregulated and 404 downregulated) at the FDR corrected q-value of 0.05.

**Figure 6.**
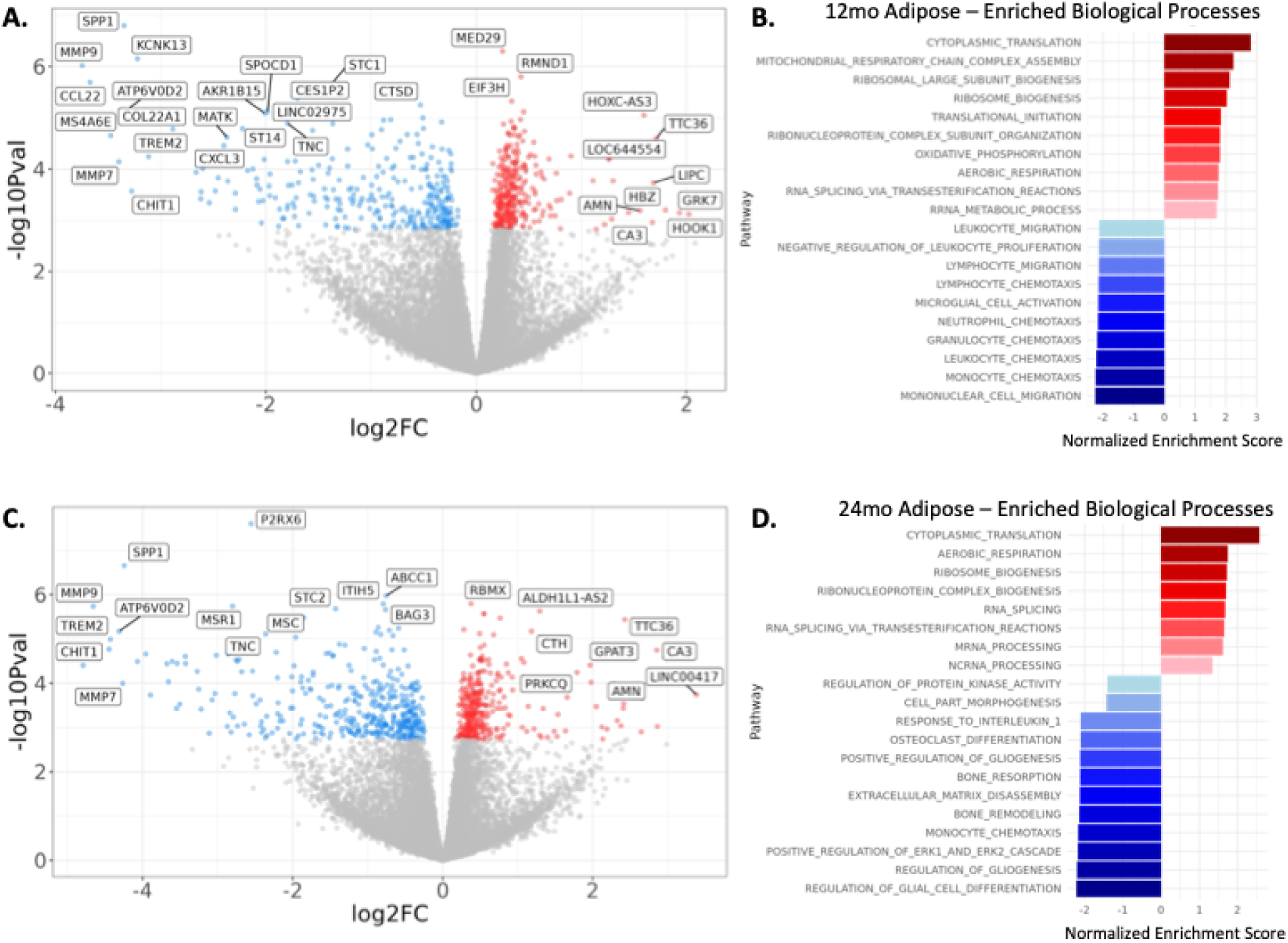
Differences in adipose mRNA expression (A and C) and enriched biological processes (B and D) between CR and AL groups at the 12-month and 24-month timepoints. For Panels A and C, x-axis shows log2 fold-change differences between groups. The y-axis shows -log10 p-value. Blue points show genes with false-discovery rate q-values less than 0.05 were downregulated in CR participants. Red points show genes with false-discovery rate q-values less than 0.05 were upregulated in CR participants. Gray points show genes with q-values equal or greater than 0.05. Using a false-discovery cutoff of q=0.05, a total of 605 genes (from 19471 genes total) were differentially-expressed between CR and AL treatment groups at the 12-month timepoint (n=34, Panel A) and 734 genes were differentially-expressed between CR and AL treatment groups at the 24-month timepoint (n=18, Panel C). A dispersed but otherwise arbitrary selection of 31 differentially-expressed genes labeled by gene symbol are shown for the 12-month treatment effect. A dispersed but otherwise arbitrary selection of 23 differentially-expressed genes labeled by gene symbol are shown for the 24-month treatment effect. Panels B and D show the top-10 upregulated (red) and down-regulated (blue) biological pathways from Gene Ontology based on overlapping results from both fast GSEA and GAGE gene set enrichment. The full set of enriched pathways for 12-month and 24-month follow-ups are in Tables S1 and S2, respectively.

For the 12-month follow-up, pathway analysis of differentially-expressed genes identified 241 enriched biological pathways (12 upregulated, 229 downregulated). Upregulated pathways included those involved in mitochondrial function, enhanced protein synthesis through ribosomal biogenesis, and RNA processing. These pathways are consistent with improved cellular energy metabolism, reduction of oxidative stress, and improved cellular repair and gene regulation, supporting longevity promoting tissue health and function through improved cellular maintenance and repair. Downregulated pathways included those involved in immune system activation and inflammatory responses, cellular and ion homeostasis, and endocytosis and cellular response to lipids. These pathways are consistent with decreases in pro-inflammatory responses, and improved handling of ions and lipids, resulting in a reduction in stress-response pathways and inflammation thought to be central to the aging process.

Moreover, these findings are consistent with a previous adipose RNAseq analysis from a subset of CALERIE participants^23^. Spadaro et al reported a similar induction of genes related to mitochondrial function and a suppression of genes related to inflammation^23^. Together, our current data, in corroboration with what has been previously reported, support the idea that caloric restriction appears to slow several hallmarks of aging in adipose tissue, including the loss of mitochondrial function and chronic inflammation^24^. A complete gene set enrichment for the 12-month follow-up is provided in **Table S4**. The top-10 upregulated and top-10 downregulated pathways for 12-month follow-up are illustrated in **Figure 6B**.

For the 24-month follow-up, pathway analysis of differentially-expressed genes identified 155 enriched biological pathway (8 upregulated, 147 downregulated). Upregulated pathways included aerobic respiration, enhanced ribosome production and protein synthesis, and gene expression regulation. These pathways are consistent with sustained improvements in cellular metabolism, repair, and gene regulation detected in adipose mRNA during the 12-month follow-up. Downregulated pathways at the 24-month follow-up also included many of the same pathways identified at the 12-month follow-up; those involved in immune system activation and inflammatory responses, cellular and ion homeostasis, and cellular metabolism. Additional pathways at the 24-month follow-up were involved in development and morphogenesis pathways, suggesting an additional benefit of CR on tissue composition and differentiation in the form of tissue maintenance. A complete gene set enrichment for the 24-month follow-up is provided in **Table S5**. The top-10 upregulated and top-10 downregulated pathways for 24-month follow-up are illustrated in **Figure 6D**.

To facilitate the rapid appraisal of the underlying data structure of smRNA and mRNA for plasma, adipose, and muscle samples, we generated a series of summary datasets for transcripts passing quality control and their distributions across each timepoint (**Supplemental Data Files 25-44**).

## Discussion

CALERIE-2™ is the first ever Randomized Control Trial (RCT) of long-term CR in healthy, non-obese humans. The CALERIE™ Genomic Data Resource reported here represents the first such resource for a trail of an intervention hypothesized to modulate biological processes of aging. Below, we briefly outline applications of this resource to advance the frontiers of translational geroscience.

First, the CALERIE™ Genomic Data Resource provides opportunities to compare impacts of novel drug and behavioral interventions designed to modulate aging biology with a gold-standard intervention well-established in the animal literature. Admittedly, CALERIE™ is an imperfect implementation of CR compared to the controlled laboratory settings of model organisms. Nevertheless, CALERIE™ provides a baseline against which alternative, potentially more scalable geroprotective interventions can be compared. The molecular functions, biological processes, and cellular components generated from genomic datasets provide a ‘common language’ for comparisons with other intervention and observational studies in humans^25^. For example, the CALERIE™ Genomic Data Resource will facilitate comparison with a growing list of behavioral and pharmacological intervention trials now underway, many of which have specified genomic variables as endpoints^26^. For example, blood DNA methylation (DNAm) are being generated in a range of intervention trials, including trials of diet and exercise^27–30^, supplements, including Vitamin D^31^ and nicotinamide mononucleotide^32^, and pharmacological interventions, including Dasatinib, Quercetin, and Fisetin^33^. RNA from blood, as well as RNA and DNAm from adipose and muscle are also available for a range of diet and exercise-related interventions^34–39^. As genomic data will become available for an increasing range of trials, the value of the CALERIE™ Genomic Data Resource will continue to grow.

Second, the CALERIE™ Genomic Data Resource will allow researchers to compare the molecular effects of CR on human subjects with the molecular responses of in vitro and in vivo model systems^40–42^. This complementary approach allows researchers to validate findings from rigorous human trails like CALERIE™ in laboratory settings and vice versa, and can provide insights into phylogenetically-conserved molecular responses to CR that can be used as benchmarks for interventions on biological aging. In one example, researchers examined the effect of CR in mice on muscle satellite cell expansion through the binding of circulating plasminogen to the Plg-RKT receptor. The role of plasminogen on muscle satellite cell expansion in mice was validated in humans using measures of blood plasminogen, a DNAm surrogate marker of plasminogen inhibitor PAI-1^43^, and muscle biopsies from the CALERIE™ trail^44^.

Through the recapitulation of findings in mice and humans, the researchers were able to demonstrate the mediating role of the plasminogen system on muscle stem cell regeneration in response to CR. The parallel analysis of human and in vivo or in vitro models has been successfully employed in the study of cancer and other diseases^45,46^ and is likely to play a key role in the future of translational geroscience.

Third, the CALERIE™ Genomic Data Resource provides opportunities to evaluate the sensitivity of proposed omics-based indices of biological aging to CR intervention. There are now abundant data to establish that intervention in the CALERIE™ trial improved a range of health parameters for the treatment group^9–11,47,48^. These data provide a background against which to evaluate proposed novel indices of biological aging, as we did in our prior analysis of epigenetic clocks^18^. The CALERIE™ Genomic Data Resource will provide a platform for analysis of the many new blood-based clocks recently published and others still forthcoming^49–52^ as well as non-omics-based measures of biological aging^18,53,54^. Furthermore, DNAm and RNA from muscle and adipose data from the CALERIE™ Genomic Data Resource will provide researchers with the rare opportunity to study the relationship between new and existing blood-based measures and molecular responses to CR in other tissues.

Fourth, the CALERIE™ Genomic Data Resource will provide developers with a powerful setting in which to develop, implement, and validate new and existing multi-omics tools^55–57^. Multi-omics approaches leverage complementary information provided by different molecular data types to gain a more comprehensive understanding of complex biological systems and processes. Multi-omics tools use similarities, network structures, correlations, and probabilistic relationships across molecular datasets to generate novel insights not evident from analysis of individual omics data^58^. Most multi-omics tools have been developed and implemented in cancer research, where they are used for studying disease biology, disease subtyping, and in the development of diagnostic and prognostic biomarkers^58^. The application of such multi-omics tools in the CALERIE™ Genomic Data Resource could be used to better understand the complex biology of CR in living humans, the heterogeneity of responses to CR, or pharmacological targets for CR mimicking drugs.

Finally, the CALERIE™ Genomic Data Resource can provide researchers with preliminary data for planning interventions on biological aging. For example, CALERIE™ results for current epigenetic clocks^18^ highlight the need for large sample sizes in trials planning to use these measures as outcomes. CALERIE™ data can similarly illuminate sample size planning around other genomic measures of aging.

In conclusion, the CALERIE™ Genomic Data Resource is a new platform to elucidate the biology of caloric restriction and its effects on biological processes of aging. This resource can also contribute to the discovery and validation of aging biomarkers and provide a reference resource for new geroscience clinical trials. It is our hope that publication of omics datasets for geroscience trials will become standard practice, advancing open science within geroscience research and enabling rapid comparative and integrative analyses to accelerate translation of therapies to extend healthspan.

## Methods

### Study Protocol

CALERIE Phase 2 was a multi-center, randomized controlled trial conducted at three clinical centers in the United States^8^ (ClinicalTrials.gov Identifier: <u>NCT00427193</u>). It aimed to evaluate the time-course effects of 25% CR (that is, intake 25% below the individual’s baseline level) over a 2-yr period in healthy adults (men aged 21–50Lyr, premenopausal women aged 21–47Lyr) with BMI in the normal weight or slightly overweight range (BMI 22.0–27.9LkgLm^−2^). The study protocol was approved by Institutional Review Boards at three clinical centers (Washington University School of Medicine, St Louis, MO, USA; Pennington Biomedical Research Center, Baton Rouge, LA, USA; Tufts University, Boston, MA, USA) and the coordinating center at Duke University (Durham, NC, USA). All study participants provided written, informed consent. Nongenomic data were obtained from the CALERIE Biorepository (https://calerie.duke.edu/apply-samples-and-data-analysis).

### Sample Collection

#### Blood Collection

Approximate 70ml of blood was collected from participants after an overnight fast (12 hours) and immediately cryopreserved in liquid nitrogen and stored at −80°C from the time of acquisition until further processing.

#### Muscle biopsies

Skeletal muscle biopsies of the vastus lateralis (VL) muscle were obtained using a 25-gauge, 2 inch needle as described in Bergstrom^59^. Participants were fasting overnight (12 hours) at the time of biosample collection. Briefly, the biopsy site was treated with local anesthesia. A small incision was made using a scalpel and the biopsy needle was inserted.L Approximately 50-100 mg of muscle tissue was extracted and placed in a cryovial and immediately flash frozen in liquid nitrogen. Samples were stored at −80°C until processing.L

#### Adipose biopsies

Abdominal subcutaneous adipose tissue biopsies were obtained using 3 or 4mm x 15cm liposuction needles. Participants were fasting overnight (12 hours) at the time of biosample collection. Briefly, an area approximately one hand-width lateral to the umbilicus was sterilized and local anesthesia obtained using 2% lidocaine. After making a small insicion with a scapel, the liposuction needle was inserted and suction was applied to the attached syringe to withdraw approximately 900mg of tissue. Tissue was rinsed with phosphate buffered saline, weighed, placed in a cryovial, and immediately flash frozen in liquid nitrogen. Samples were stored at −80°C until processing.

#### Genotyping

The CALERIE™ genotype dataset was produced by the Kobor Lab at the University of British-Columbia and the Genomics Analysis Shared Resource at Duke University. DNA was extracted from n=217 baseline blood samples obtained from the CALERIE™ Biorepository at the University of Vermont. Participants were fasting overnight (12 hours) at the time of biosample collection. Genotyping was conducted using the Illumina Global Screening Array-24 v3.0 (GSA) BeadChips containing 654,027 markers, with ∼30,000 add-on markers from Infinium PsychArray-24 focused content panel (Illumina, San Diego, CA). Briefly, 200ng DNA was processed and hybridized to the GSA chips according to the manufacturer’s instructions, and scanned using the Illumina iScan platform. Genotypes were called for SNPs matched to dbSNP v151 ^60^ and for which valid calls were made in >98% of participants using GenomeStudio v2.0 (Illumina, San Diego, CA). Additional SNPs were imputed using the IMPUTE2 software suite G3^61^ and the 1000 Genomes Phase 3 reference panel^62^. Pre-phasing and imputation were conducted by breaking the genome into 5 megabase fragments. The final SNP database consisted of directly genotyped SNPs and SNPs imputed with a >90% probability of a specific genotype for which call rates were >95% and minor allele frequencies were >1%.

#### SNP-principal components

The top 20 principal components were calculated on the autosomal SNPs using PLINK v1.90b3.36 with default parameters^63^.

### DNA methylation

#### Preprocessing, Normalization, and Quality Control

##### Blood

DNA methylation profiling for blood was conducted in the Kobor Lab (https://cmmt.ubc.ca/kobor-lab/) from whole-blood DNA stored at −80 degrees Fahrenheit. Briefly, 750ng of DNA was extracted from whole blood and bisulfite converted using the EZ DNA Methylation kit (Zymo Research, Irvine, CA, USA). Quantity and quality of the bisulfite-converted DNA using NanoDrop spectrophotometry. Methylation was measured from 160ng of bisulfite-converted DNA using the Illumina EPIC Beadchip (Illumina Inc, San Diego, CA, USA).

QC and normalization were performed using methylumi^64^(v2.32.0) and the Bioconductor (v 2.46.0)^65^ package from the R statistical programming environment (v 3.6.3). Probes with detection p-values >0.05 were coded as missing; probes missing in >5% of samples were removed from the dataset (final probe n=828,613 CpGs). Normalization to eliminate systematic dye bias in the 2-channel probes was carried out using the methylumi default method.

##### Muscle

DNA methylation profiling for muscle was conducted in the UCLA UCLA Neuroscience Genomics Core (UNGC) from skeletal muscle DNA extracted in the VB Kraus lab at Duke University and stored at −80 degrees Celcius. Briefly, 250 ng of DNA was extracted from muscle tissue and bisulfite converted using the EZ DNA Methylation kit (Zymo Research, Irvine, CA, USA).

Methylation was measured from 250 ng of bisulfite-converted DNA using the Illumina EPIC Beadchip (Illumina Inc, San Diego, CA, USA). Quality control and preprocessing for muscle tissue DNAm was carried out from IDATs in the Geroscience Core in the Butler Columbia Aging Center, and performed in the R computing environment using the same methods described for blood DNA methylation. The final probe number for muscle DNA methylation was n=861779.

##### Adipose

DNA methylation profiling for adipose was conducted in the UCLA Neuroscience Genomics Core (UNGC) from adipose tissue DNA extracted in the VB Kraus lab at Duke University and stored at −80 degrees Celcius. Briefly, 250 ng of DNA was extracted from adipose tissue and bisulfite converted using the EZ DNA Methylation kit (Zymo Research, Irvine, CA, USA). Methylation was measured from 250 ng of bisulfite-converted DNA using the Illumina EPIC Beadchip (Illumina Inc, San Diego, CA, USA). Quality control and preprocessing for adipose tissue DNAm was carried out from IDATs in the Geroscience Core in the Butler Columbia Aging Center, and performed in the R computing environment using the same methods described for blood and muscle DNA methylation. The final probe number for adipose DNA methylation was n=861714.

#### Derived Variables

##### DNAm control-probe principal components (PCs)

For blood, muscle, and adipose, we conducted principal component analysis of EPIC-array control-probe beta values to compute controls for technical variability across the samples. To do so, we loaded raw idats, normalized using *normalizeMethyLumiSet*, and extracted normalization quality control probes. Principal components (PCs) were extracted from normalization probes, and standard deviations of these PCs were standardized by squaring and dividing by the sum of their squares. PCs that explained 90% of the variance in normalization probes were retained.

##### Immune cell proportion estimates

For blood, proportions of CD8T, CD4T, natural killer cells, b cells, monoctyes, and neutrophils were estimated using the Houseman equation via the *FlowSorted.Blood.EPIC* package and standard “Blood” reference panel^66,67^(**Supplementary Data File 39**). An ‘extended’ version of *estimateCellCounts2* using *FlowSorted.BloodExtended.EPIC* using the “BloodExtended” reference panel was also used to calculate basophils, B naïve, B memory, CD4T naïve, CD4T memory, CD8T naïve, CD8T memory, eosinophils, monocytes, neutrophils, T regulatory cells and natural killer cells (**Supplementary Data File 40**). Cell estimates were estimated on a red green channel set (RGset) and noob normalized with the reference dataset as specified by the package developers^19^. For contrast between tissues, we illustrate deconvoluted cell types (adipose, epithelial, fibroblast, and immune cells) for blood, muscle, and adipose tissues using hierarchical EpiDISH^68^ in Figure S2.

##### Epigenetic clock estimates

Blood epigenetic clocks including Horvath, Hannum, PhenoAge, GrimAge were calculated using the matrix of beta values provided in the data repository and the online calculator (https://dnamage.genetics.ucla.edu/, since been superseded by https://projects.clockfoundation.org). DunedinPACE was calculated using the matrix of beta values provided in the data repository and the DunedinPACE R package (https://github.com/danbelsky/DunedinPACE). Principal components-based clocks (PC clocks) developed by Higgins-Chen and colleagues were calculated in R on noob normalized samples that passed sample quality control using previously described code and methods^69^.

### RNAseq

#### Blood

##### RNA extraction

Circulating RNA was extracted from 200 µL plasma using the Qiagen miRNeasy Serum/Plasma Advanced Kit (Catalog no. 217204). RNA samples were stored at −80°C until miRNA sequencing was performed.

##### sRNA sequencing

sRNA sequencing was performed on samples isolated from plasma (n=584; AL=36%:CR=64%). Sequencing libraries were prepared from 5µLs of miRNeasy RNA using the QiaSeq miRNA Library Auto Kit (384; cat # 331509) according to the manufacturer’s instructions. Library amplification (22 cycles) was performed on a Beckman i7 liquid handler, using QiaSeq miRNA 96 Index IL Auto A (384; cat # 331569). Library quality was evaluated using a fragment analyzer (Agilent); the library concentrations were quantified by Qubit ds DNA HS assay (ThermoFisher). Libraries were sequenced on a NovaSeq 6000 (Illumina) using S4 lanes and 75 base pair single reads.

##### sRNA processing and normalization

QIAseq smRNA sequencing FASTQ files were preprocessed using the GeneGlobe® Data Analysis platform, Legacy version 2.0. SmRNA reads were first processed by trimming at the 3’ adapter using cutadapt. Next, the trimmed reads were used to identify insert and unique molecular index (UMI) sequences. Reads with <16 base pair insert sequences or less than 10 base pair UMI sequences were discarded. Annotations of insert and UMI sequences were mapped to the human reference genome (GRCh38/hg38) using the alignment tool, bowtie2 (v2.5.1). All miRNA/piRNA mapped reads and associated UMIs were then aggregated to count unique molecules using miRbase V21 and piRNABank, respectively. Following the data processing steps, UMI counts were filtered to remove low expression smRNAs (row sum counts <50 were removed). Filtered UMI counts were then used to calculate normalization factors using the trimmed mean of M-values (TMM) method within the BioConductor edgeR (v4.2.1) package.

#### Muscle

##### RNA extraction

Flash frozen skeletal muscle samples (∼50 mg) were homogenized using a bead-based Qiagen TissueLyser II in l mL TRIzol™ Reagent for a total of 4 minutes, rotating the TissueLyser Adapter after 2 minutes (Invitrogen™ ThermoFisher #15596026). Following homogenization, RNA was extracted using a Qiagen RNeasy Mini Kit (Qiagen #74104) and stored at −80°C until mRNA sequencing.L

##### mRNA sequencing

Illumina libraries were generated with the TruSeq Stranded Total RNA Library Preparation Kit. RNA was sequenced using the *Illumina NovaSeq 6000* sequencing system with paired-end reads. The samples yielded 387 to 618 million pass filter reads with more than 86% of bases above the quality score of Q30.

##### mRNA filtering, alignment, and genome annotation

The quality ofLreads in fastq RNA-Seq files were initially assessed using *FastQC* tool (v0.11.5; https://www.bioinformatics.babraham.ac.uk/projects/fastqc/), Preseq (v2.0.3)^70^, Picard tools (v2.17.11; https://broadinstitute.github.io/picard/) and RSeQC (v2.6.4)^71^. Reads were trimmed using *bbduk* (from bbtools package (https://jgi.doe.gov/data-and-tools/software-tools/bbtools/)^72^. Following trimming, cleaned reads were examined one more time using *FastQC*.L Next, cleaned fastq files, along with reference human genome 38 and Ensembl annotation v104, were used as input for STARL (v2.7.10a), a splice-aware aligner implemented with a novel algorithm for aligning high-throughput long and short RNA-Seq data to a reference genome^73^.L The STAR aligner was run with the –*quantmode TranscriptomeSam* parameter to also generate transcriptome BAM files, and was used for RSEM analysis.LGenome BAM files were sorted and indexed using samtools.LFinally, genome BAM files wereLused as input for *featureCounts* from the Rsubread package (v2.0.1)^74^, a suitable program for counting reads for various genomic features such as genes.

##### sRNA sequencing

sRNA sequencing was performed on samples isolated from muscle (n=116; AL=38%:CR=62%) using the same protocols as for blood, described above.

##### sRNA Processing and Normalization

sRNA data were processed and normalized using the same protocols as for blood, described above.

#### Adipose

##### RNA extraction

Segments of flash frozen abdominal adipose tissue (∼50 mg each) were homogenized using a bead-based Qiagen TissueLyser II in l mL TRIzol™ Reagent for a total of 4 minutes, rotating the TissueLyser Adapter after 2 minutes (Invitrogen™ ThermoFisher #15596026). Following homogenization, RNA was extracted using the Qiagen RNeasy Mini Kit (Qiagen #74104) and stored at −80°C until smRNA sequencing.

##### mRNA sequencing

mRNA sequencing was performed on the Illumina NovaSeq 6000 Sequencer at the Duke Sequencing Core. Barcoded libraries for mRNA sequencing were produced from the Universal Plus mRNA-seq NuQuant kit (0408-A01, Tecan Genomics) according to manufacturers recommended protocol with 17 amplication cycles and 200ng of total starting RNA. The protocol was automated on a Perkin Elmer Sciclone G3 liquid handler.

##### mRNA filtering, alignment, and genome annotation

Alignment of fastq files against the GRCh38 assembly of the human genome was performed using Subread^75^ (v2.0.3) and bowtie2 (v2.5.1) on the Columbia Department of Systems Biology high performance cluster. Most samples aligned with >90% mapped reads. One sample aligned with 69.6% mapped reads, but demonstrated satisfactory base sequence quality, tile sequence quality, and per sequence quality scores. Quantification was performed using featureCounts^74^ method as implemented in Subread. A matrix of counts was constructed, with genes labeled using Entrez Gene ID numbers. Initial quality control steps and outlier detection was carried out in 5 groups of 30 samples each. Samples that passed the first screen were then combined for a final quality and outlier detection round. Quality was assessed and outliers detected by multidimensional scaling^76^ and hierarchical clustering^77^. Multidimensional scaling was performed using limma^78^ (v3.54.1). Hierarchical clustering was performed with Euclidean distance and complete linkage clustering^77^ as implemented in the Heatmap.2 command in gplots^79^. Optimal clusters determined by the gapstat^80^ method as implemented in NBClust^81^ (v3.0.1). During quality control, 2 samples – both baseline samples – were removed. One sample had too few reads, while the other was a clear outlier on multidimensional scaling plots and linkage clustering maps.

##### mRNA Differential Expression

Samples were normalized using the trimmed mean method^82^ and analyzed for two-sided differential expression with limma-voom with sample weighting to downweight the effect of sample outliers^78,83,84^. Participant was modeled as a random effect using the duplicate correlation method^85,86^. CR at follow-up (12- or 24-months) was compared to AL at follow-up (12- or 24-months), relative to CR at baseline compared to AL at baseline. Models included covariates for sex, age at baseline, BMI and RNA integrity number. Raw p-values were corrected for multiple testing using the Benjamini-Hochberg false discovery rate^87^. Limma estimates and effect-sizes are provided for 12-month and 24-month follow-up in **Table S2** and **Table S3**, respectively.

For gene set enrichment analysis, we used a custom R function that integrates outputs from the fast GSEA (FGSEA; v1.30.0) and GAGE (v2.54.0) methodologies^88,89^. We focused on GO Biological Processes (GO-BP), with c5.go.bp.v7.5.symbols.gmt (downloaded from the Molecular Signature Database, http://www.gsea-msigdb.org/gsea/msigdb/) as our pre-defined gene set. FGSEA was executed with parameters set to a minimum pathway size of 15, a maximum of 600, and 10,000 permutations. Pathways with an FDR adjusted p-value below 0.05 were retained. To enhance the robustness of our findings, GAGE analysis was also performed, filtering pathways significant in both FGSEA and GAGE. We categorized pathways as either up-regulated or down-regulated based on their normalized enrichment scores and visualized the top 10 up- and down-regulated pathways.

##### sRNA sequencing

sRNA sequencing was performed on samples isolated from muscle adipose (n=65 AL=38%:CR=62%) using the same protocols as for blood, described above.

##### sRNA processing and normalization

sRNA data were processed and normalized using the same protocols as for blood, described above.

## Data Availability

Processed data can be accessed through the Aging Research Biobank (https://agingresearchbiobank.nia.nih.gov/studies/calerie/). Data use is restricted to non-commercial use in studies to determine factors that affect age-related conditions. Original raw data may be obtained from the Belsky Lab. Code used in the production of summary data and figures are available at https://github.com/CPRyan/CALERIE_Genomic_Data_Resource.

## Acknowledgement

This research received support from US National Institute on Aging Grants R01AG061378, R33AG070455, R01AG054840, and U01AG060908 and from US National Cancer Institute Grant 5P30CA013696. DWB is a Fellow of the CIFAR CBD Network. This research utilized the FlowSorted.BloodExtended.EPIC software packages developed at Dartmouth College, which are governed by the licensing terms provided by Dartmouth Technology Transfer (https://github.com/immunomethylomics/FlowSorted.BloodExtended.EPIC/blob/main/SoftwareLicense.FlowSorted.BloodExtended.EPIC%20to%20sign.pdf).

