## Supplementary Materials for "The CALERIE™ Genomic Data Resource"

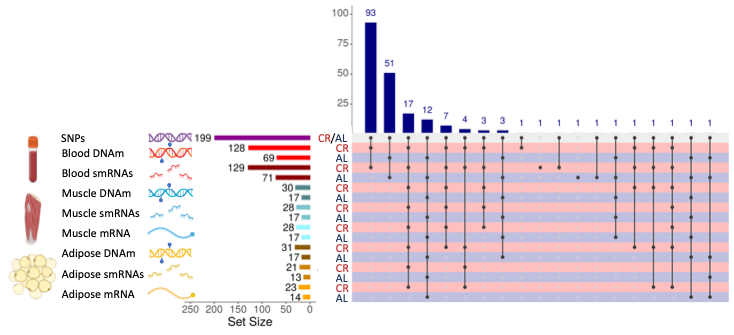

**Figure S1.** The figure shows an upset plot displaying overlap of available datasets across treatment group, tissues, molecular data type. This figure is analogous to Figure 2, Panel A in the main manuscript, with the exception that here only samples with baseline and at least one follow-up (12- or 24-months) are counted. The left-hand side shows tissue type (blood, muscle, or adipose), type of molecular data type (SNPs, DNAm, smRNAs, or mRNA), set size for each tissue and data type combination in number of unique individuals, and group (CR or AL). The bottom right-hand side shows points and connecting lines indicating overlapping intersections across tissues and data types color coded by treatment group (CR = red, AL = navy). The top right hand side shows a barchart indicating sample sizes for the overlapping intersections of tissue and datatypes. Each tissue and molecular data type combination is linked to a corresponding color scheme as follows: genomic variation (purple DNA), blood DNA methylation (red DNA with lollipops), blood small RNAs (smRNAs; dark red RNA fragments), muscle DNA methylation (blue DNA with lollipops), muscle smRNAs (blue RNA fragments), muscle mRNA (blue single RNA strand), adipose DNA methylation (yellow DNA with lollipops), adipose smRNAs (yellow RNA fragments), adipose mRNA (yellow single RNA strand).

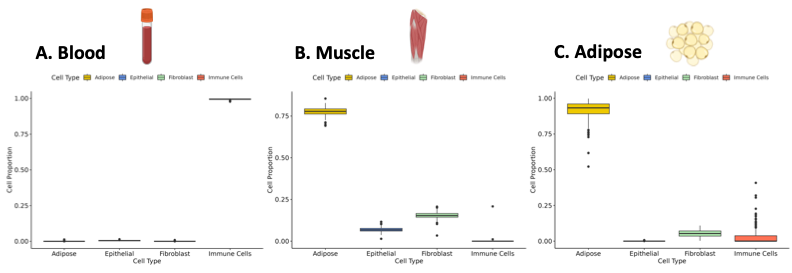

**Figure S2.** The figure shows estimated adipose, epithelial, fibroblast, and immune cell proportions for blood (Panel A), muscle (Panel B), and adipose (Panel C) tissue samples. Estimated cell proportions are based on DNA methylation and the hierarchical EpiDISH deconvolution method described in Zheng et al. 2018; Epigenomics.

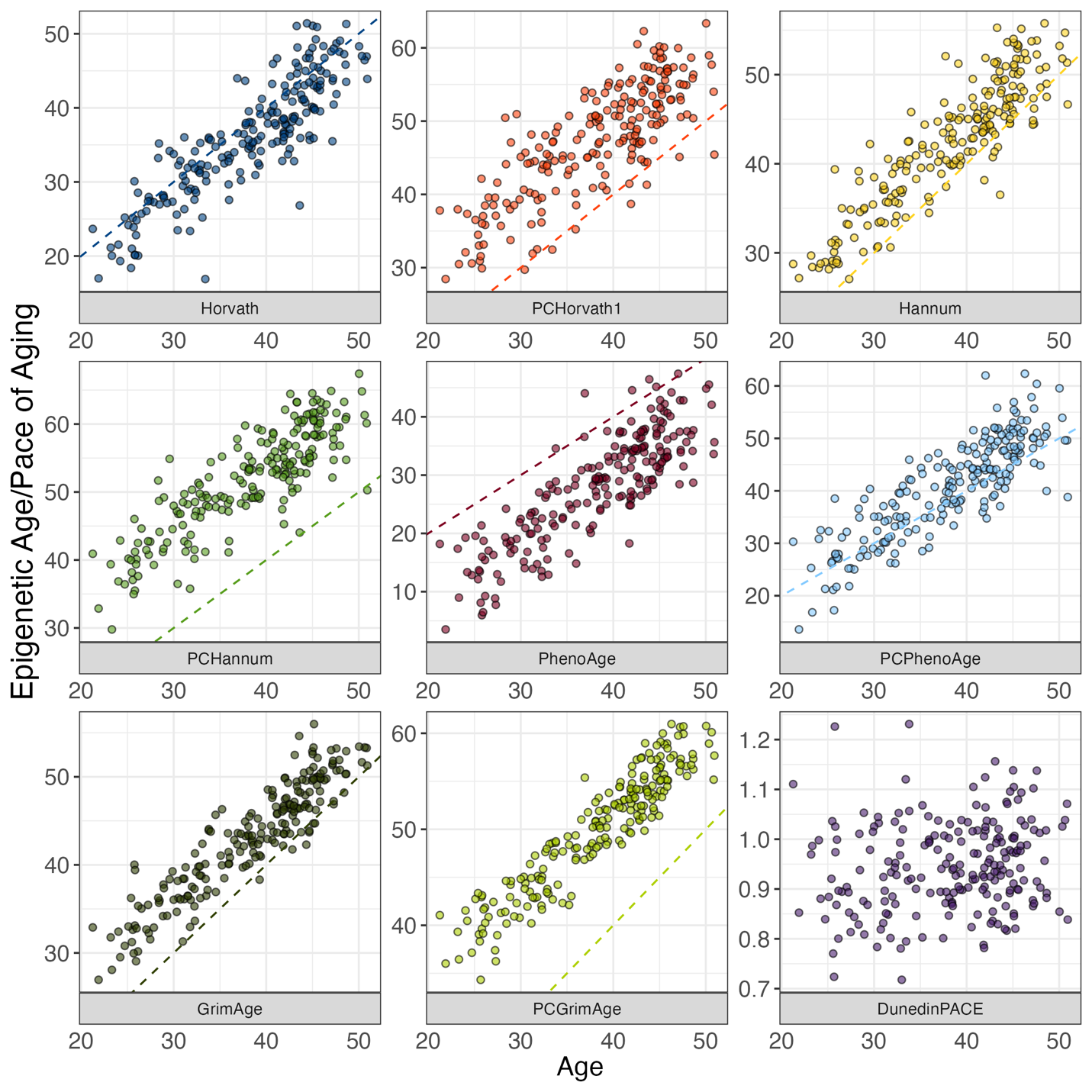
**Figure S3.** Associations of DNA methylation measures of aging with chronological age. The figure shows DNA methylation measures of aging (Y-axis) against chronological age (X-axis) for n=212 men and women at pre-intervention baseline. The dashed colored line on each facet is the line of identity (intercept=0, slope=1), indicating where predicted epigenetic age or pace of aging would equal chronological age. Correlations with chronological age are as follows: Horvath Clock r=0.87, PC Horvath Clock r=0.84, Hannum Clock r=0.92, PC Hannum Clock r=0.88, Skin & Blood Clock r=0.94, PC Skin & Blood Clock r=0.80, PhenoAge Clock r=0.84, PC PhenoAge Clock r=0.85, GrimAge Clock r=0.93, PC GrimAge Clock r=0.92, DunedinPACE r=0.15.

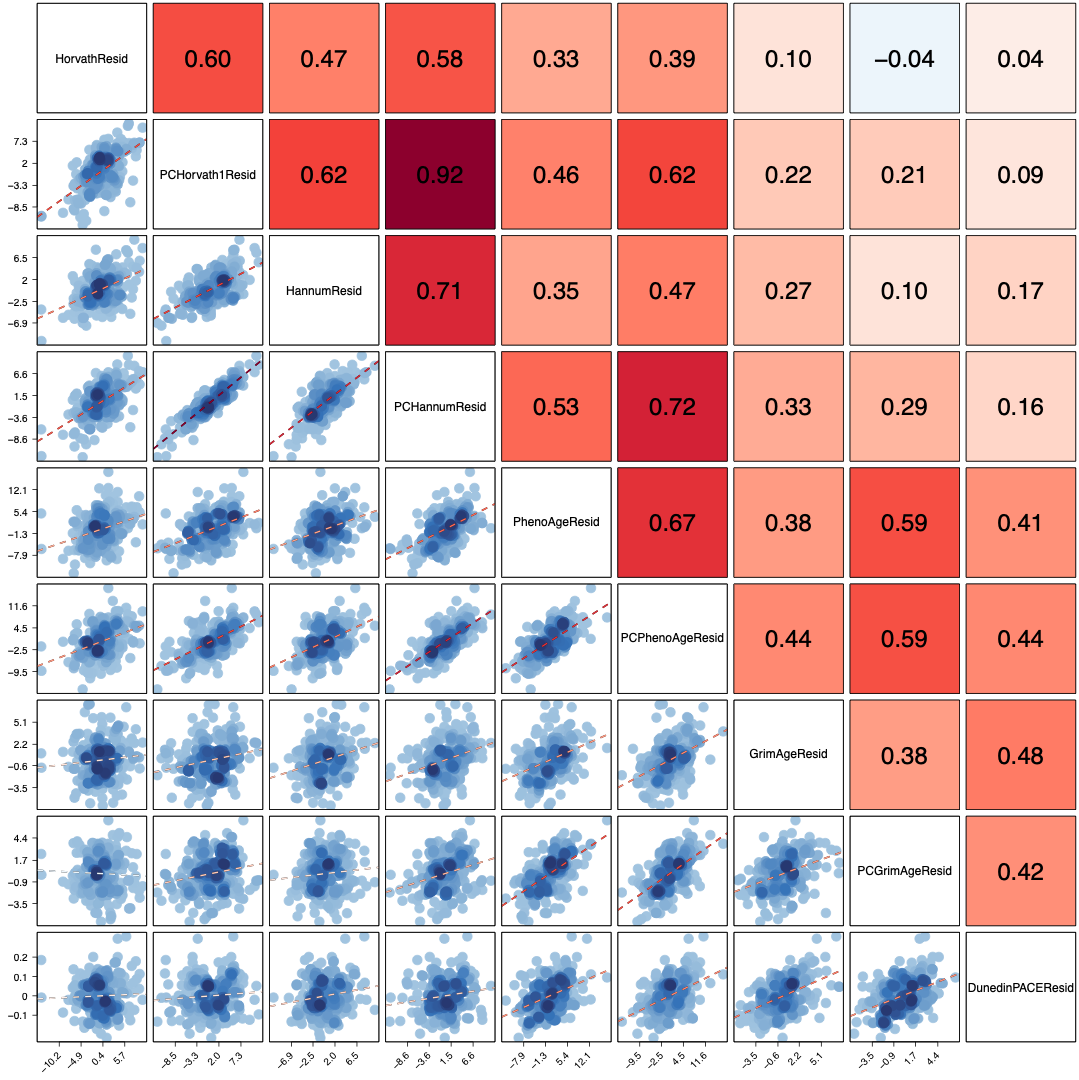

**Figure S4. Associations of age-residualized DNA methylation measures of aging with each other.** The figure shows correlations between age-residualized DNA methylation measures of aging for n=212 men and women at pre-intervention baseline with each other. The dashed red line on each facet is the fitted regression slopes. Pearson correlations between DNA methylation measures of aging are shown on the upper diagonal facets, with the shade of the facet indicating the strength of the correlation.
